# Levels and patterns of genetic diversity differ between two closely related endemic *Arabidopsis* species

**DOI:** 10.1101/048785

**Authors:** Julie Jacquemin, Nora Hohmann, Matteo Buti, Alberto Selvaggi, Thomas Müller, Marcus A. Koch, Karl J. Schmid

## Abstract

Theory predicts that a small effective population size leads to slower accumulation of mutations, increased levels of genetic drift and reduction in the efficiency of natural selection. Therefore endemic species should harbor low levels of genetic diversity and exhibit a reduced ability of adaptation to environmental changes. *Arabidopsis pedemontana* and *Arabidopsis cebennensis*, two endemic species from Italy and France respectively, provide an excellent model to study the adaptive potential of species with small distribution ranges. To evaluate the genome-wide levels and patterns of genetic variation, effective population size and demographic history of both species, we genotyped 53 *A. pedemontana* and 28 *A. cebennensis* individuals across the entire species ranges with Genotyping-by-Sequencing. SNPs data confirmed a low genetic diversity for *A. pedemontana* although its effective population size is relatively high. Only a weak population structure was observed over the small distribution range of *A. pedemontana*, resulting from an isolation-by-distance pattern of gene flow. In contrary, *A. cebennensis* individuals clustered in three populations according to their geographic distribution. Despite this and a larger distribution, the overall genetic diversity was even lower for *A. cebennensis* than for *A. pedemontana.* A demographic analysis demonstrated that both endemics have undergone a strong population size decline in the past, without recovery. The more drastic decline observed in *A. cebennensis* partially explains the very small effective population size observed in the present population. In light of these results, we discuss the adaptive potential of these endemic species in the context of rapid climate change.

## Introduction

A main concern in the current debate on ecological effects of climate change is whether populations and species can adapt fast enough to keep up with the rapid rate of environmental changes (Salamin *et al*. 2010). Mitigation of biodiversity loss requires knowledge on ecological and evolutionary responses of populations to habitat changes, and on the conditions that allow a recovery of declining populations (Gonzalez *et al*. 2012). Plant endemic species are an important component of biodiversity, particularly in biodiversity hotspots, where they make up a large proportion of the local flora (Myers *et al*. 2000). With a limited geographic distribution that is often tied to specific habitats, endemic species are more vulnerable to environmental changes as they frequently depend on the existence of particular biotic and abiotic interactions (Thomas *et al*. 2004). For this reason endemic biodiversity should be a central focus of conservation efforts. Because of their limited distribution range, endemic species tend to have small census population sizes and smaller effective population sizes, N_e_ (Freville *et al*. 2001; Strasburg *et al*. 2011). Theory predicts that a small effective population size leads to slower accumulation of mutations, increased levels of genetic drift and reduction in the efficiency of natural selection (reviewed by Ellstrand & Elam 1993). According to the neutral theory of molecular evolution, genetic diversity levels at neutral sites reflect a balance between the mutational input per generation and the loss of genetic variation due to genetic drift (Kimura 1983). All else being equal, species with smaller population sizes should thus harbor lower levels of neutral genetic diversity (Ramos-Onsins 2004; Leimu *et al*. 2006; Leffler *et al*. 2012). The chance of new, potentially advantageous mutations appearing is also reduced compared to larger populations. Additionally, hard selective sweeps from new mutations and soft selective sweeps from standing variation are less likely, reducing the rate of adaptation (Lanfear *et al*. 2014). Furthermore, genetic drift may lead to the fixation of mildly deleterious mutations which are not efficiently removed because of weak purifying selection (Kimura 1983; Cao *et al*. 2011; Xue *et al*. 2015), leading potentially to a mutational meltdown (Lynch *et al*. 1993). Consequently, the direct and indirect effects of a small effective population size have a strong influence on the evolutionary dynamics and the adaptive potential of a species (Charlesworth 2009). Continuous adaptation in changing environments requires sufficient and appropriate genetic variation that provides extreme genotypes capable of surviving intense stress conditions and allow the population persistence (Reed *et al*. 2011; Bell 2013). Low levels of genetic diversity, reduced average fitness and limited adaptive potential have been indeed observed for plant populations with small population size (Pluess & Stöcklin 2004; Hensen & Oberprieler 2005; Leimu *et al*. 2006; Michalski & Durka 2007; Leimu & Fischer 2008). A meta-analysis demonstrated that genetic erosion significantly contributes to the extinction risk of plant species, beside short term demographic and ecological processes (Spielman *et al*. 2004). To evaluate the adaptive potential and demographic responses of endemic populations in a rapidly changing environment, it is therefore necessary to estimate levels of genetic diversity and to infer the evolutionary processes (genetic drift, gene flow, mutation, mating system) that have shaped the pattern of genetic variation (Reed *et al*. 2011).

In the genus *Arabidopsis*, the two endemic species *Arabidopsis cebennensis* (DC.) and *Arabidopsis pedemontana* (Boiss.) provide an excellent model of species with restricted ranges. *Arabidopsis thaliana* and its close relatives have become model plant species in population genetics and the data accumulated for the whole genus enable an interspecific comparative approach of genetic diversity levels (Mitchell-Olds & Schmitt 2006; Clauss & Koch 2006). The whole genus reflects the diversity of plant distribution ranges: while *A. thaliana* is distributed world-wide, *A. cebennensis* and *A. pedemontana* have the smallest distribution range in the genus (O’Kane & Al-Shehbaz 1997; Koch *et al*. 2008). *Arabidopsis cebennensis* is restricted to the mountainous Massif Central region in Southern France at elevations ranging from 900 to 1500 m a.s.l. Highly disjunct populations occur over an area of about 11,000 km^2^ in the Cevennes, Cantal, Aveyron and Ardèche regions. *Arabidopsis pedemontana* occupies a much smaller distribution range of 50 km^2^ in the Piedmont region of the northwestern Italian Alps, at altitude ranging from 1,300 to 2,200 m a.s.l. All known populations of this species occur on the two sides of a single mountain ridge located between the valleys Po and Pellice. *Arabidopsis pedemontana* is included in the red list of Italian and Piedmont Floras under the category “critically endangered' according to the IUCN definition (The IUCN Species Survival Commission 2004), whereas *A. cebennensis* is not protected but occurs mostly in national and regional protected reserve areas. The two species are closely related to each other and their common ancestor is genetically distinct from other lineages in the genus (Koch & Matschinger 2007; Hohmann *et al*. 2014). They are perennial diploids and presumably self-incompatible, with a strong tendency for vegetative reproduction by clonal growth (Hohmann *et al*. 2014). Both species occupy a very specific niche in riverine habitats (streams and waterfalls), with a semi-continental mountainous climate (cold winters). Adaptation to this specific habitat led to a very specialized ecology, and phenotypically differentiate them from their relatives which are found in different habitats like meadows or rocky outcrops. Previous studies revealed a highly reduced diversity for chloroplast DNA, ribosomal DNA and microsatellite loci in *A. cebennensis* and *A. pedemontana* compared to other *Arabidopsis* species (Koch & Matschinger 2007; Hohmann *et al*. 2014). Due to this reduced level of genetic variation, these surveys have not provided much information on genetic structure. Whole genome approaches are likely more promising.

The present work aims to reveal the extent and structure of genetic variation in the two endemic *A. cebennensis* and *A. pedemontana* on a genome-wide scale. We used Genotyping-By-Sequencing (GBS, Elshire *et al*. 2011) to obtain genome-wide SNPs for estimating population genetic parameters. GBS and other reduced-representation sequencing approaches (Davey *et al*. 2011) have rapidly become important tools for the study of genetic diversity, adaptation and conservation (Narum *et al*. 2013; Huang *et al*. 2014; Gonçalves da Silva *et al*. 2015; Xue *et al*. 2015). As the genomes of both endemic species have not yet been sequenced, we used *A. lyrata*, which is the most closely related species with a sequenced genome, as a reference for mapping and SNP calling. Additionally we analyzed the structure of genetic variation at the population level based on a set of microsatellite loci and plastid DNA sequences from the *trnLF* region, which have been used previously to characterize *A. cebennensis* and *A. pedemontana* genetic diversity within the whole genus *Arabidopsis* (Hohmann *et al*. 2014).

Based on an extensive sampling representing 90% and 40% of all known locations for *A. pedemontana* and *A. cebennensis*, respectively, we characterized the genetic diversity and the contemporary effective population size of both endemic species over their entire distribution range. We also investigated how their demographic history may have led to the observed levels of genetic variation, and tested whether it is consistent with models of declining population sizes. In light of these new results, we discuss the adaptive potential of the two endemic species in the context of the probable environmental changes to come.

## Materials and Methods

More detailed information on the materials and methods is available in Text S1.

### Plant material

All samples were collected between 2010 and 2014 in France and Italy (Fig. 1, Supporting information, Hohman *et al*. 2014). For GBS, genomic DNA was extracted using a modified CTAB protocol (Saghai-Maroof *et al*. 1984). For samples collected more recently, DNA was extracted with the Genomic Micro AX Blood Gravity kit (A&A Biotechnology, Gdynia, Poland). DNA concentration and quality was checked with agarose gel electrophoresis and Qubit 2.0 Fluorometer (Life Technologies). In the GBS analysis, 53 *A. pedemontana* and 28 *A. cebennensis* samples were used.

**Figure 1.**
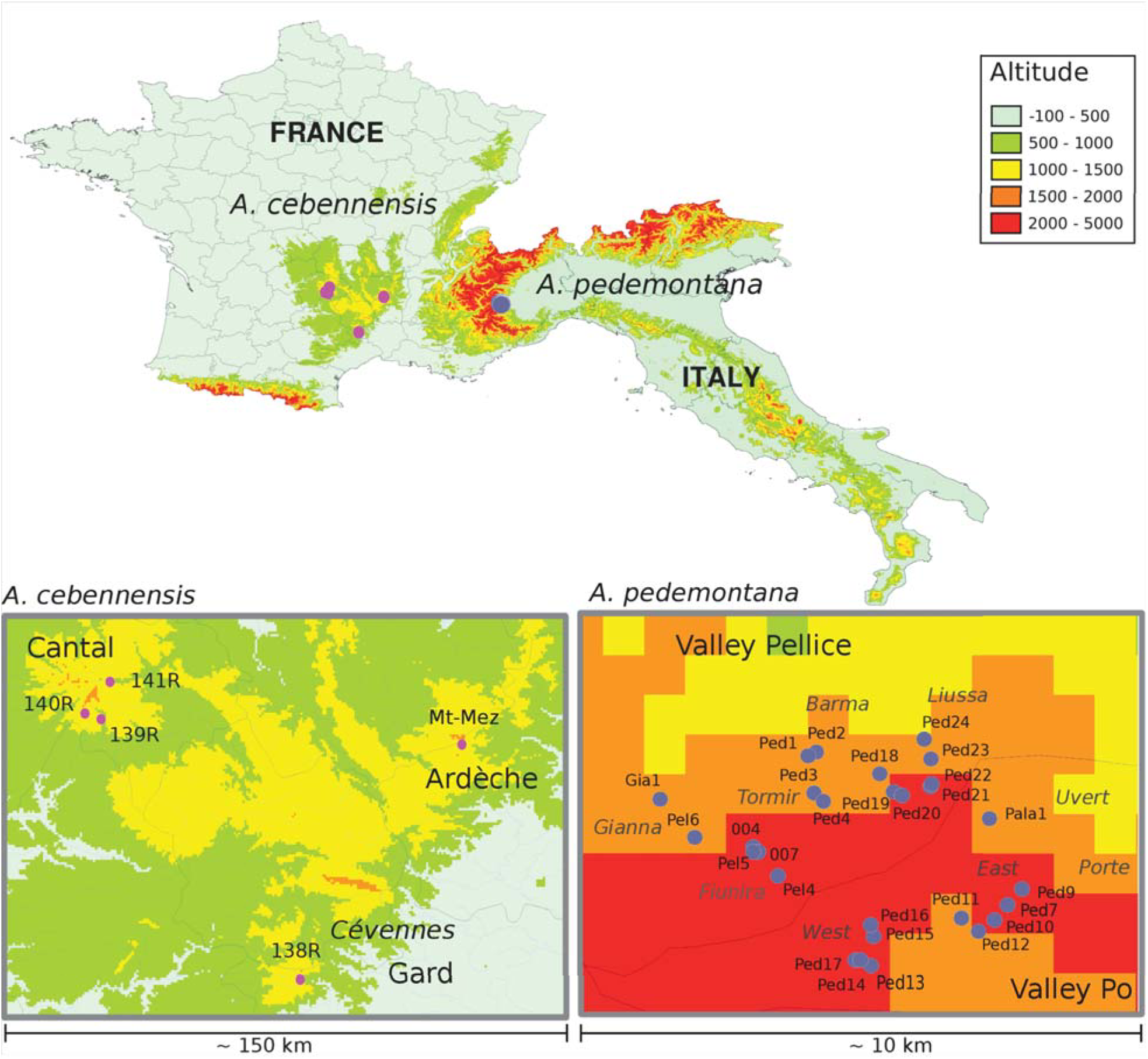
Distribution range and altitude of the populations and individuals sampled for *A. cebennensis* and *A. pedemontana*.

### Genotyping-by-Sequencing (GBS)

Double-digest GBS libraries were constructed based on the design of (Poland *et al*. 2012) with ApeKI as rare-cutting and HindIII as common restriction enzyme. Adapters and primers (Metabion) were taken from (Elshire *et al*. 2011). Adapters were modified to introduce 58 different barcodes and sticky ends corresponding to ApeKI and HindIII cut sites (Fig. S1). Five μl of each samples with different barcodes were pooled after adapter ligation. Three pools were prepared, one with 24 *A. pedemontana* individuals, one with 29 *A. pedemontana* individuals, and one pool including all 28 *A. cebennensis* samples. The pooled samples were purified with QIAquick PCR purification kit (Qiagen, Hilden/Germany) and PCR amplified. The GBS libraries were run on BluePippin (Sage science) for selection of DNA fragments ranging between 200-350 bp. Each of the three pools was sequenced on one lane of the same flowcell, on an Illumina HiScanSQ with single end sequencing and 105 cycles. With a genome size of around 250 Mbp for both species (Lysak *et al*. 2009; Hohmann *et al*. 2014) the targeted coverage per site was between 20 and 30x.

### Sequence data analysis and SNP calling

Raw reads were processed with custom Python scripts, bwa (Li & Durbin 2009) and FastQC >(http://www.bioinformatics.babraham.ac.uk/projects/fastqc/). All reads with ambiguous ‘N’ nucleotides and reads with low quality values (< 90% bases with Q>20) were discarded. Read length after barcode and end-trimming was 90 bp (Supporting information). The pre-processed reads were aligned to the genome of *Arabidopsis lyrata* strain MN47 (Hu *et al*. 2011) with pBWA (Peters *et al*. 2012) with 10 mismatches allowed. Biallelic SNPs were called with SAMtools (Li *et al*. 2009) and custom Python scripts from aligned reads with a minimum mapping quality of 1. The vcf file was parsed to filter out SNPs with a coverage of at least 30x and at most 3000x, where by at least five reads had to confirm the variant nucleotide. Five *A. pedemontana* and seven *A. cebennensis* individuals with less than 5,000 SNPs were excluded to reduce the proportion of missing values (Supporting information). The SNPs were further filtered using vcftools (Danecek *et al*. 2011) to include only intra-specific *A. pedemontana* and *A. cebennensis* polymorphic sites (Table S2). Two additional files were created including only SNPs with data for at least 50% and 70% of the sampled individuals (Table S2). SNP calling was conducted additionally for ten different subsets of 21 randomly chosen *A. pedemontana* individuals, in order to compare the number of SNPs observed for *A. cebennensis* and *A. pedemontana* when working with the same sample size. The total number of sites (polymorphic and non-polymorphic) and variants (SNPs and indels) were also calculated, applying the same filters described above, allowing us to determine the percentage of polymorphic loci for both species.

### Structure and genetic diversity analysis

Population structure was inferred using SNPs with data for at least 50% of the sampled individuals. The optimal number of clusters was obtained using ADMIXTURE (Alexander *et al*. 2009)with the cross-validation procedure (‐‐cv) with K ranging from 1 to 9. Ten iterations with different seed values were completed and compared for homogeneity. Discriminant Analysis of Principal Components (DAPC; Jombart *et al*. 2010) was conducted using the adegenet R package, first using the function find.clusters to determine the optimal number of clusters (K), with K ≤ 10. 25 and 60 principal components (PCs) were kept to explain the variance in *A. cebennensis* and *A. pedemontana* respectively. Phylogenetic networks were generated in Splitstree (Huson & Bryant 2006) after vcf files were converted into IUPAC coded (Cornish-Bowden 1985) FASTA files. Distances were calculated using the Uncorrected_P method, ignoring ambiguous states, and the network was generated using the Neighbor-net distances transformation (Bryant & Moulton 2004) with 100 bootstrap replicates. A network was also constructed for the two species and including the reference *A. lyrata* as outgroup, using SNPs called between the three species. To investigate the extent to which the genetic distance between populations was affected by spatial structure (dispersal limitation), and to test the isolation-by-distance (IBD) hypothesis (Slatkin 1993), a Redundancy Analysis (RDA) was performed (Borcard *et al*. 1992) using the R package ade4 (Text S1). The presence of migration events between *A. cebennensis* large populations was tested by running TreeMix v1.12 (Pickrell & Pritchard 2012) with *A. pedemontana* as outgroup, and four migration events allowed. One thousand bootstrap replicates were generated by resampling blocks of 500 SNPs.

All population genetic parameters were calculated from SNPs with data for at least 70% of the sampled individuals. The percentage of missing data was calculated using the R package adegenet 1.4-2 (Jombart & Ahmed 2011). Nucleotide diversity (π) was calculated for each SNP and then averaged over the total number of sites to obtain an average nucleotide diversity per bp, using the formula in Begun *et al*. (2007). Watterson's estimator (θ_w_) was calculated to compare the number of segregating sites between the populations. Nei's gene diversity (or proportion of heterozygosity expected, H_exp_) as well as the proportion of heterozygosity observed (H_obs_) were calculated using adegenet. The sum of expected heterozygosity per polymorphic sites was divided by the total number of sites to calculate the mean expected heterozygosity. F_st_ was calculated per site with the R package pegas 0.6 (Paradis 2010), using the formula of Weir & Cockerham (1984). Mean values of F_st_ were calculated over all sites. The partitioning of genetic variability among and within populations in *A. cebennensis* was analyzed with an AMOVA in Arlequin v.3.5, using 10,000 permutations and pairwise difference as distance calculation method.

### Demographic analysis

The long-term demographic history of the two species was investigated with δaδi v1.7.0 (Gutenkunst *et al*. 2009). We examined five one-dimension models for each species independently, one neutral and four population size change models (Fig. S2). The standard neutral model (SNM) assumes a constant population size (ancestral population size N_a_). For the simple exponential size change model 1a, the population size has changed instantaneously at time *T* in the past to yield a present population size of N. Model 1b assumes that the population size has changed exponentially since time *T* in the past, leading to a contemporary population size *N*. Model 1c assumes that the species have first experienced an instantaneous size change, yielding a population size of *N*_0_, followed by an exponential population size change, going from *N*_0_ to *N* in time *T*. In Model 1d, we expand Model 1a by incorporating a second instantaneous size change event. At time *T* + *T*_0_ in the past, the population goes through a size change of depth *N*_0_, and then recovers to relative size *N*. The last two models allow us test the probability of a past bottleneck event followed by either recovery or further decrease of the population size. Additionally we tested three two-populations models in which *A. cebennensis* and *A. pedemontana* diverged from an ancestral population (Fig. S2). In model 2a, at *T*_*pc*_ × 2*N*_*a*_ generations ago, the two species split, yielding a stable contemporary population size *N*_*c*_ for *A. cebennensis* and *N*_p_ for *A. pedemontana*. In model 2b, the split is followed by exponential size change of both new species, going from *N*_*c*0_/*N*_*p*0_ to *N*_*c*_/*N*_*p*_ during the time *T*_*pc*_ since the divergence. In model 2c, we incorporated a bottleneck event in the ancestral equilibrium population before the divergence of the species. The depth and duration of the bottleneck are denoted by *N*_0_ and *T*_0_ respectively. The Python script implementing the models is provided as Supporting information.

The models were fitted to the SNPs dataset with data for at least 70% of the sampled individuals. The data joint site frequency spectrum (SFS) was estimated by δaδi, and projected down to 15 individuals for *A. cebennensis*, nine for *A. cebennensis* Cantal population alone, and 40 individuals for *A. pedemontana*, which provided 6,583, 3,981, and 10,422 SNPs for the singlepopulation models. For the two-population models, the data was projected down to a sample of 14 and 30 individuals for *A. cebennensis* and *A. pedemontana* respectively, which provided a large number of segregating sites (10,442). The SFS were folded. Parameters for each model were optimized with the upper and lower bounds, arbitrary start values and grid sizes indicated in Supporting information. Multiple optimizations runs were evaluated for each model to ensure the convergence of the optimized parameters. To estimate parameter's confidence intervals (CIs), 100 replicate pseudo-data sets were generated by bootstraping the SNP data by 1 Mb regions on the *A. lyrata* scaffolds to account for putative linkage between the SNPs. CIs were estimated using the SFS of the bootstraped data sets. The log composite likelihood of each model was calculated using δaδi's *ll_multinom* function and the likelihood ratio test was used to test if the differences in the likelihood values between the models were significant.

### Microsatellite and plastid DNA sequence analysis

In a previous work, 148 *A. cebennensis* individuals from ten populations and 40 *A. pedemontana* individuals from nine populations were genotyped with seven microsatellite loci and sequences from the *trnLF* region (Hohmann *et al*. 2014). Detailed experimental protocols are available in this study. We carried out a population structure analysis of the two endemic species based on this dataset. Geographic locations of the populations analyzed, total numbers of alleles and mean number of allele per locus for the seven microsatellite loci are indicated in Fig. S3 and Supporting information. Microsatellite genotypes were analyzed using STRUCTURE v.2.3.4 (Pritchard *et al*. 2000; Hubisz *et al*. 2009). Ten replicates were run for each K-value and a burn-in-period of 1 × 105 and 2 × 105 iterations was used. The option 'admixture model' was used in combination with 'correlated allele frequencies'. The estimation of the optimal K number of populations (ranging from 1 to 5) was calculated using the R-script Structure-sum (Ehrich 2006). Input files for CLUMPP were generated with STRUCTURE HARVESTER (Earl & VonHoldt 2012), alignments of replicate runs were conducted in CLUMPP (Jakobsson & Rosenberg 2007), and the mean of 10 runs were visualized.

## Results

### GBS and polymorphisms

Forty-eight *A. pedemontana* individuals from 29 locations and 21 *A. cebennensis* individuals from five locations were included in the final analysis. Only results based on SNPs with data for at least 70% of the sampled individuals are presented, except if noted otherwise. The average read coverage per site over all individuals was 465 for *A. cebennensis* and 1,124 for *A. pedemontana*. The number of SNPs and percentage of polymorphic loci over all sites were more than two times higher in *A. pedemontana* than in *A. cebennensis* (Table 1). When the SNP calling was conducted on ten different subsets of 21 randomly sampled *A. pedemontana* individuals, the resulting average number of SNPs was 11,858 SNPs (Table S3), which is only ∼1,000 less than the number of SNPs observed for the total sample of *A. pedemontana*. This shows that the different levels of polymorphism in *A. cebennensis* and *A. pedemontana* are not caused by contrasting sampling sizes but truly indicate a higher level of polymorphism in *A. pedemontana*.

**Table 1.** GBS analysis output: Number of total and polymorphic sites. Only SNPs with data for at least 70% of the sampled individuals are included.

|  | *A. cebennensis* | *A. pedemontana* |
| --- | --- | --- |
| Total sites | 874,955 | 739,868 |
| SNPs | 6,583 | 12,909 |
| % polymorphic sites | 0.7 | 1.74 |
| Average read depth per SNPs | 465 | 1,124 |
| % missing data | 13.7 | 7.9 |

The Neighbor-net network constructed for the two species and the outgroup *A. lyrata* shows a clear distinction between them, with high bootstrap support for the branches leading to each species (Fig. S4). *Arabidopsis pedemontana* and *A. cebennensis* are closer to each other in the network. 667 common SNPs called independently in *A. cebennensis* (10%) and *A. pedemontana* (5%) have shared alleles, indicating a high level of shared variation between the two sibling species.

### Population structure

Based on the geographic distribution of the samples (Fig. 1), our initial hypotheses were: 1) Genetic variation in *A. cebennensis* should primarily cluster according to the three large regions where this taxon is observed in the Massif central; 2) Distinct patterns of variation should be observed for *A. pedemontana* between the two valleys where the species occurs, as the mountain range constitutes a natural barrier to dispersion. In the ADMIXTURE analysis the lowest crossvalidation value was always found when assuming K=2 for *A. cebennensis*, followed closely by K=3 (Fig. S5). With K=2, individuals from Cantal are separated from Ardèche and Cévennes individuals. The clustering for K=3 (Fig. 2) corresponds exactly to the origin of the individuals, from the Ardèche, Cévennes and Cantal regions. For K=4, individuals from Cantal were further split into two groups, with location 139R separated from locations 140R and 141R. For *A. cebennensis*, six PCs and two discriminant functions were retained when describing clusters with DAPC, which supported the K=3 clustering. The Neighbor-Net network of *A. cebennensis* also separated the Ardèche, Cantal and Cévennes populations with strong bootstrap support (Fig. 2). Within the Cantal population, individuals from site 139R clustered apart with strong support while individuals from 140R and 141R sites were mixed. We used TreeMix to test for gene flow between the three allopatric populations of *A. cebennensis*. In the resulting maximum likelihood tree, Ardèche and Cévennes populations are grouped together with a strong support (bootstrap 99.8%; Fig. S6). Only a single migration event, from Ardèche to Cantal, was inferred, but it shows a low bootstrap value (52%) and low average weight over all bootstrap replicates (13%). This indicates no or very limited gene flow between the three *A. cebennensis* populations. The genetic assignment by STRUCTURE based on microsatellite data also resulted in an optimal value of K=3, reflecting the distribution of sampled populations in their respective geographic regions (Fig. S3), although with some level of admixture, specially between Ardèche and Cévennes individuals. However, one very small population from the Cantal region (145R) is most closely associated with the genetic cluster found in the Cévennes region. This population is located at an artificial creek close to a forest road side. Considering that also its chloroplast genome type is identical to the Cévennes region (Fig. S3), it is more likely that we observed here a very rare case of recent long-distance dispersal, maybe promoted by humans. Strong phylogeographic signal in *A. cebennensis* is highly supported by the distribution of maternally inherited plastid DNA types (*trnLF* loci). In the three major regions, distinct types prevail and we did not find variation within a population. The Redundancy analysis (RDA) on genetic diversity with spatial variables explained 92.5% of the total variance in genetic diversity. However the spatial variables (distance) did not have a significant contribution (p>0.05), rejecting isolation by distance as responsible for the strong structure observed in *A. cebennensis*.

**Figure 2.**
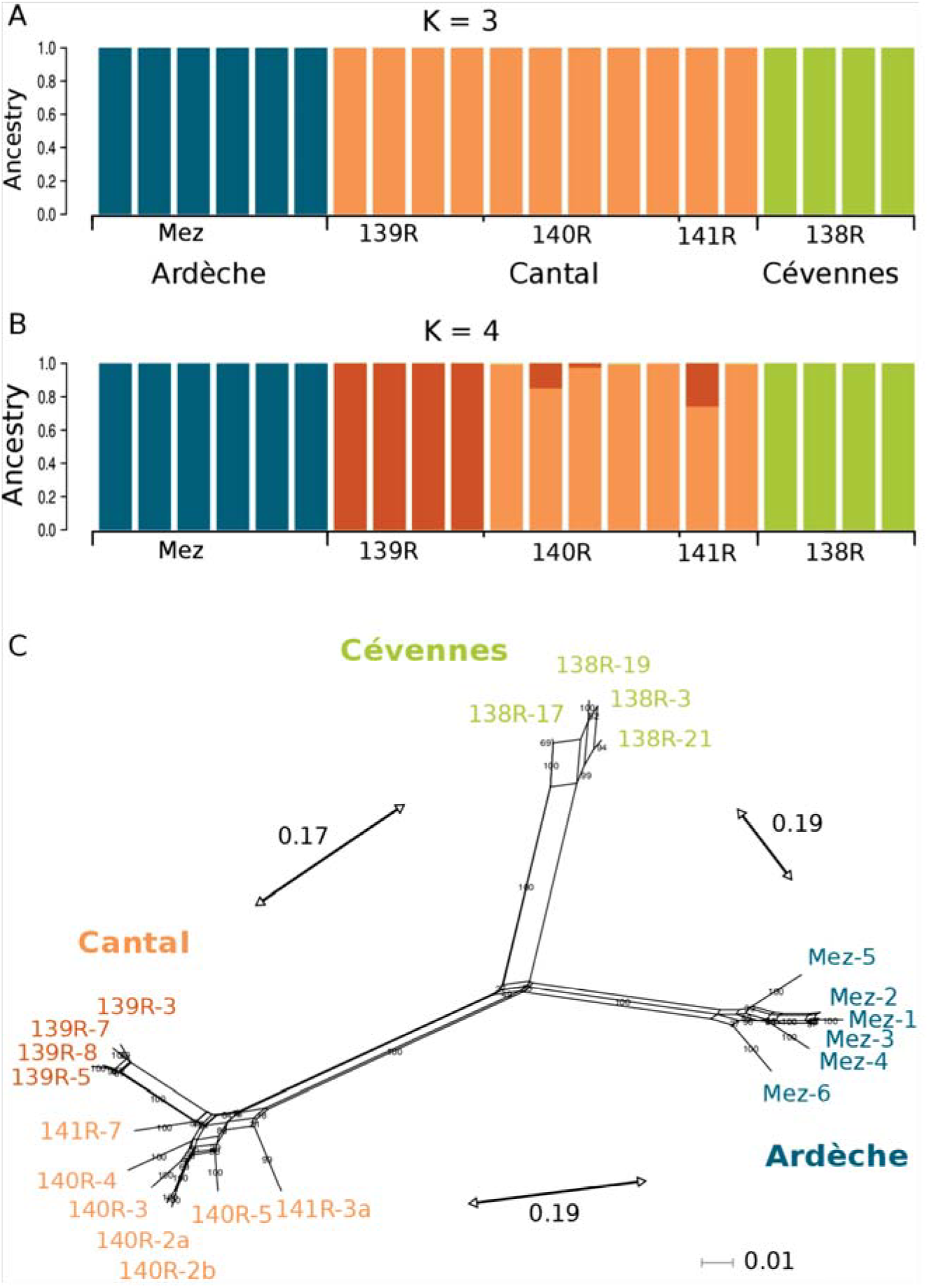
Population structure analysis of *A. cebennensis:* A) ADMIXTURE plots for number of clusters K=3 and K=4, B) Neighbor-net phylogenetic network. Bootstrap values are indicated on the network branches. F_st_ values between the three large populations are indicated along the arrows.

For *A. pedemontana*, the ADMIXTURE analysis did not suggest population structure as the cross-validation value slowly increased for K=1 to K=9 (Fig. S5). Inspection of population assignment for K=2 did not confirm our hypothesis of a differentiation of *A. pedemontana* individuals between the two valleys. Instead, individuals from Vallone di Fiunira (Fiunira) clustered as one genotype while individuals from Comba della Gianna (Giana) presented an admixture of both detected genotypes (Table S1). When describing clusters with DAPC, 15 PCs and 1 discriminant functions were retained. The DAPC analysis confirmed the results of the ADMIXTURE analysis (Fig. 3). Phylogenetic analysis of *A. pedemontana* confirmed a weak population structure with a star-shaped network, with low support for internal edges and long terminal branches (Fig. 3). However, clusters could still be observed that corresponded to the geographic distribution of the samples. For example, all individuals from Valle Po were clustered,with subgroups for the East and West part of the valley with a good bootstrap support. All individuals from Fiunira and Gianna were also grouped together, although this large cluster was not strongly differentiated from the remaining individuals and showed a weak bootstrap support. Within this group, the samples from the site 007 appeared very closely related to each other, showing the homogeneity of individuals in one location. The same was observed for the site 004. Although the microsatellite sampling for *A. pedemontana* was less complete, it contained one ipopulation that was not sampled with GBS (114; Supporting information). The genetic assignment tests for microsatellite data did not find a significant K exceeding 1, confirming the GBS results. If “LocPriors” were set and varying from 2 to 9, a significant K=2 was found,separating population 114 from the other investigated populations (Fig. S3). It is consistent with I the fact that geographic distances are correlated to genetic differentiation in *A. pedemontana*, as population 114 was the only one sampled in Valley Po for this dataset. However, with the more uniformly distributed samples in the GBS analysis, no higher-level structure is apparent between valley Po and valley Pellice. RDA analysis on genetic diversity with spatial variables explained a significant contribution of 13.5% of the total variance in genetic diversity found in populations (p<0.05). Hence dispersal limitation due to geographic distances (isolation-by-distance model) appears to be responsible for the weak structure observed in *A. pedemontana*. The level of plastid DNA variation is very low as only the ancestral type A was found (Fig. S3).

**Figure 3.**
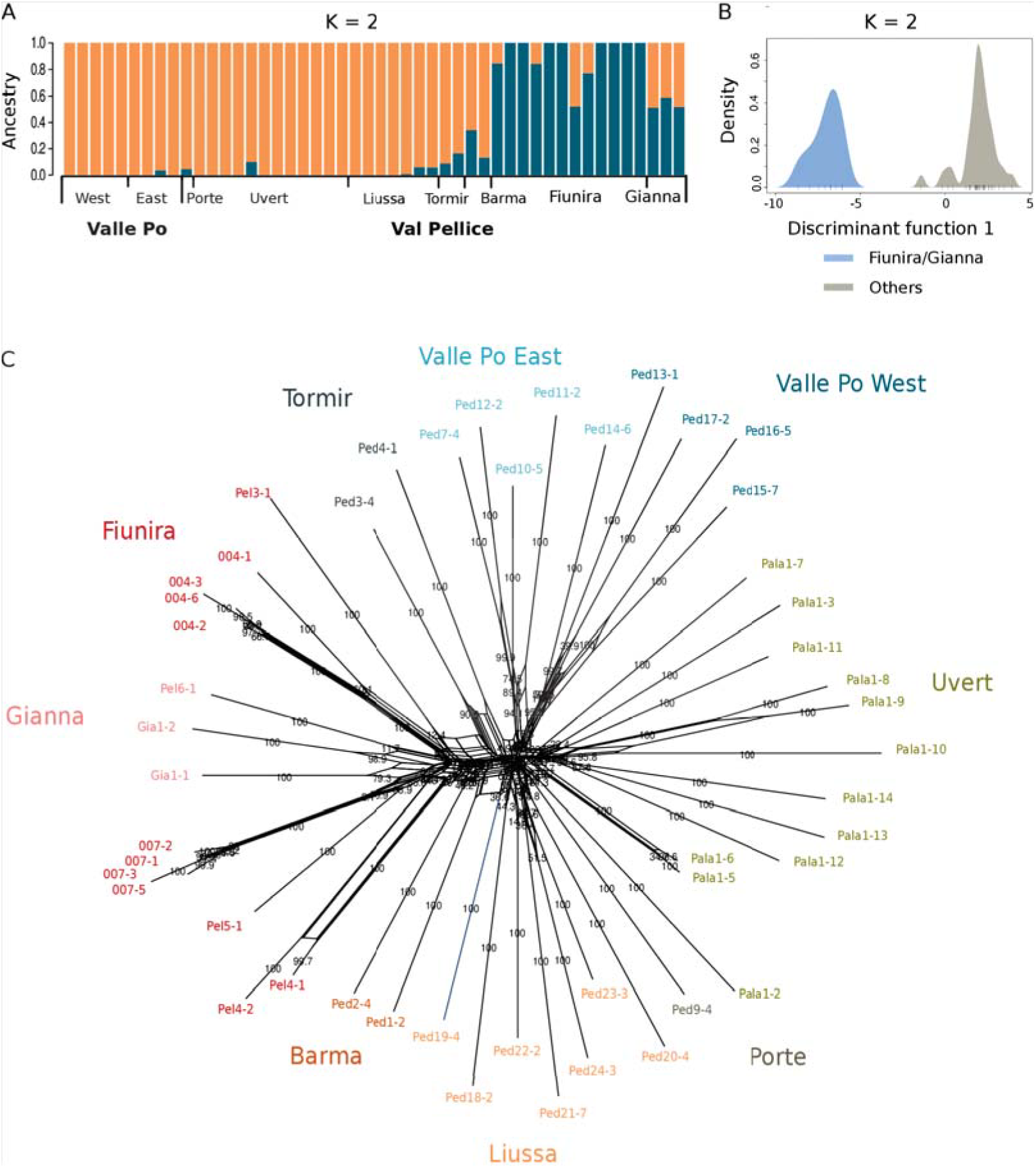
Population structure analysis of *A. pedemontana:* A) ADMIXTURE plots for number of clusters K=2, B) Plot of DAPC showing the first principal component of the analysis with K=2, with individuals (vertical bars) and groups (colored courbs) plotted on the plane, C) Neighbor-net phylogenetic network. Bootstrap values are indicated on the network branches.

### F_st_-based analysis of genetic differentiation

The mean pairwise genetic distance based on F_st_ values between *A. cebennensis* populations were similar (~0.17-0.19 ± 0.007), with the greatest distance observed between Ardèche and Cantal (Fig. 2). F_st_ values were high compared to the distance between the two species (0.33 ± 0.005). Within the Cantal population, the subpopulation 139R and 140/141R displayed a much lower mean F_st_ over sites (0.05 ± 0.005). *Arabidopsis pedemontana* individuals from Fiunira/Gianna clustered together showed a mean F_st_ of 0.04 ± 0.001 with the rest of *A. pedemontana* samples, and such a weak differentiation did not support this clustering. The distributions of F_st_ values between the two species and between the three subpopulations of *A. cebennensis* were homogeneous, with high and low F_st_ observed equally all along *A. lyrata* scaffolds used as reference genome (Fig. S7).

### Intraspecific genetic diversity

The nucleotide diversity per base pair averaged on total number of sites (π) was calculated for both species (Table 2). π was 0.0026 for *A. cebennensis* and 0.0040 for *A. pedemontana*. The distribution of π per SNP was homogeneous along *A. lyrata* scaffolds for both species (see Manhattan plots in Fig. S8), indicating that the genetic variability within each species is distributed uniformly along chromosomes. The three *A. cebennensis* populations displayed similar values of π (Table 2). θ_w_ values were almost identical to π (Table 2). *Arabidopsis cebennensis* also displayed a lower mean expected heterozygosity (H_exp_) than *A. pedemontana* (Table 2). The mean observed heterozygosity (H_obs_) calculated were only slightly lower than the expected values. These results showed a higher diversity overall in *A. pedemontana* sampled populations compared to *A. cebennensis*. In *A. cebennensis*, 39% of the genetic variability in the species is explained by the population structure (among populations), while the rest is explained by individuals within populations.

**Table 2.**
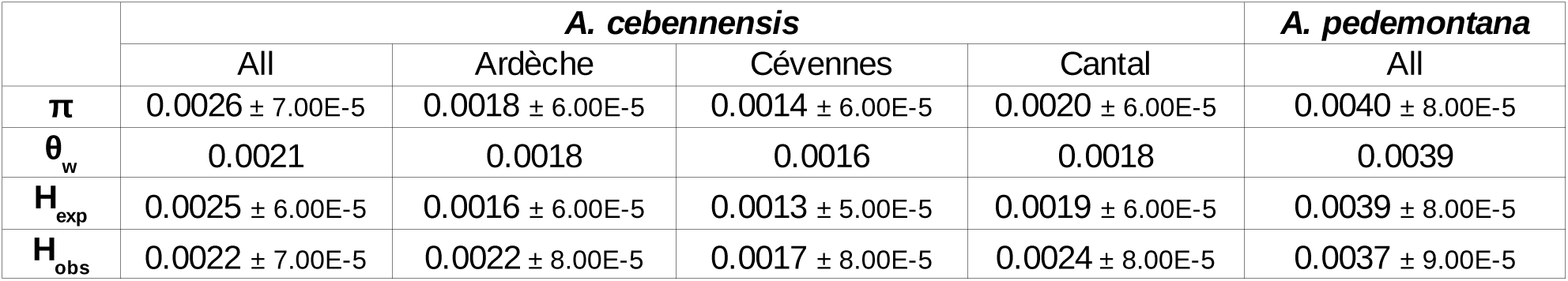
Intra-specific genetic diversity parameters

### Species demographic history

To infer the history of the two species, we tested five single-population demographic models using the joint site frequency spectrum (SFS) and the maximum likelihood methods implemented in δaδi. Considering the western alpine and sub-alpine distribution of the two species, and the fact that they occur at the highest possible elevation of the surrounding regions, we assumed they diverged during a glaciation area, and survived only in refugia in higher altitude during warming period, where they were already adapted to the colder conditions. Such a scenario implies a strong population decline of the species somewhere in their past. For each species, we fitted models 1a to 1d to test for different scenarios of population size change and compared the results to the standard neutral model (SNM) (Fig. S2). We were unsuccessful in testing models incorporating a split of *A. cebennensis* in three sub-populations, due to the sampling size and the quantity of segregating sites obtained for each of these populations. The observed minor allele frequency spectra of *A. cebennensis* and *A. pedemontana* were quite different. In *A. pedemontana*, the declining function (larger number of rarer alleles) is consistent with mutation-drift equilibrium in a stable population (log composite likelihood value LL = −334.4 for SNM). *Arabidopsis cebennensis* SFS was right-shifted (biased towards intermediate frequency alleles), deviating from the neutral expectation (LL = −1229.8 for SNM). This was explained by the strong population structure in *A. cebennensis*. The δaδi analyses suggest that the instantaneous population size change model 1a and the exponential size change model 1b are better than the SNM based on LL values and fitting plots (Fig. 4 and Fig. S2). The two models 1a and 1b fitted the observed data significantly better than the dual size change model 1c (with an error alpha= 5%, Adjusted D-statistic = −0.79 for *A. cebennensis* and −3.75 for *A. pedemontana*, p-value = 1). Model 1d fitted the observed data poorly (Fig. S2).

**Figure 4.**
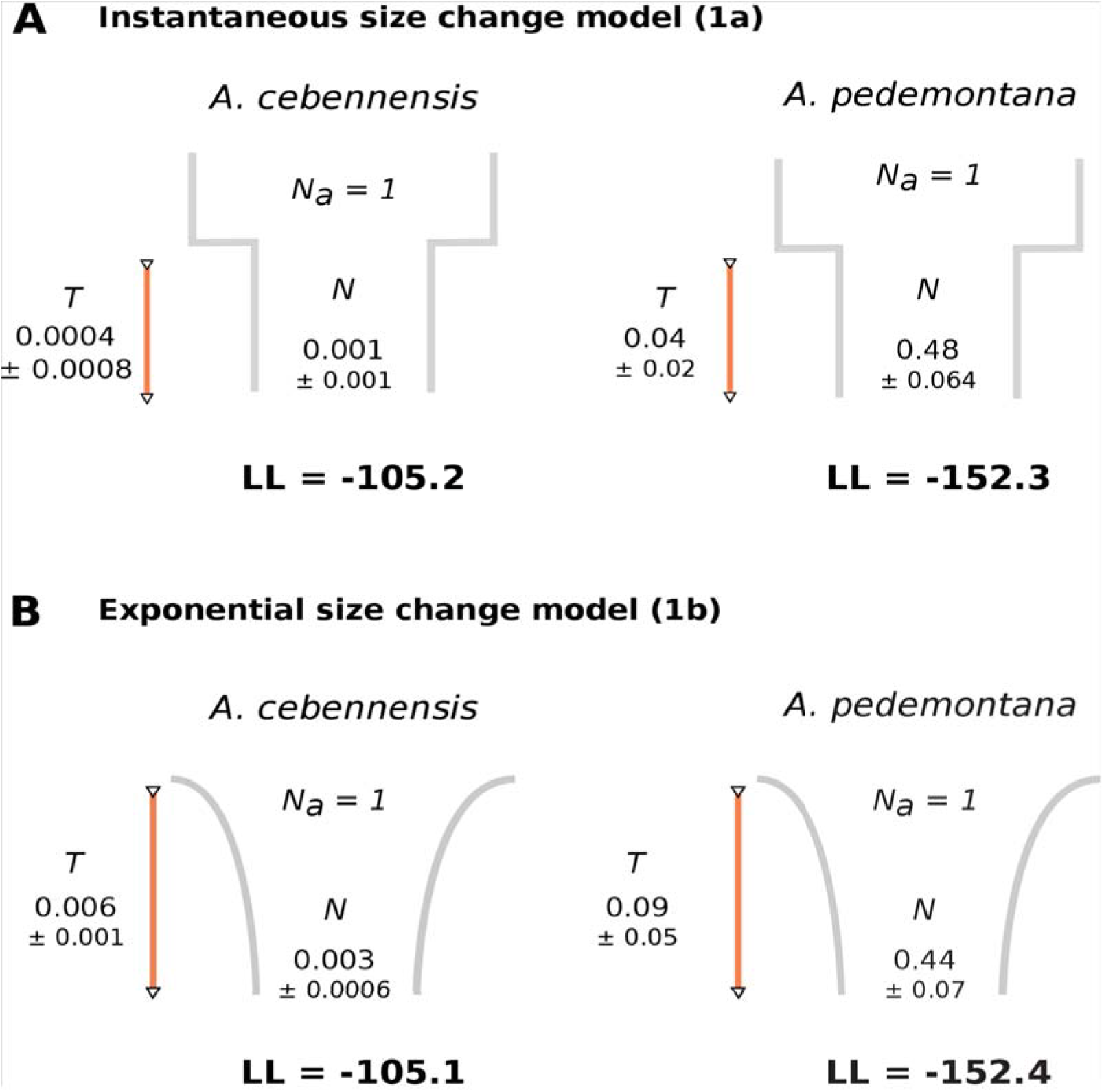
Representation of demographic models 1a and 1b and their optimized parameters fitted to observed data in *A. cebennensis* and *A. pedemontana*. The log likelihood values (LL) are indicated for each model.

Equivalent fits were observed for models 1a and 1b, which could not be compared with a likelihood ratio test (LRT) as they were not nested. Both suggested a strong population decline for the two species (Fig. 4) and indicated a deeper reduction and smaller relative population size for *A. cebennensis:* the effective population size of this species has been reduced to approximately 0.1% of its ancestral population size N_a_ (Fig. 4). In contrast, the *A. pedemontana* population shrunk only to half of its N_a_. Assuming an average generation time of two years and an overall mutation rate of 7 × 10^-9^ base substitutions per site per generation (Ossowski *et al*. 2010) we estimated the modern effective population size of *A. cebennensis* to be ≈ 140 [0-287] from model 1a or ≈ 450 [365-548] from model 1b. The effective population size for *A. pedemontana* was ≈ 55,000 [47,455-62,056] from model 1a and ≈ 50,000 [42,377-58,411] from model 1b. Our estimates placed the drastic population size decrease in *A. cebennensis* either ≈ 230 [0-689] years ago (model 1a) or ≈ 3,700 [3,044-4,262] years ago (model 1b). The decrease of *A. pedemontana* population was estimated either ≈ 18,000 [9,126-27,378] (model 1a) or ≈ 41,000 years ago (model 1b).

As the population structure observed in *A. cebennensis* was not modeled in the analysis, it could explain the non-optimal fitting between the models SFS and *A. cebennensis* observed SFS, the larger correlated residuals for *A. cebennensis*, and the strikingly low population size estimated for this species. To evaluate the effect of strong population structure on model fitting, we additionally fitted all single-population models to the observed data of the Cantal population separately. This population, for which we have the largest sample size (nine individuals), provides a representative dataset for the whole species without effect of structure. However the SFS observed for the Cantal population was also biased toward intermediate allele frequencies (Fig. S2) and the single-population models did not fit the Cantal data better than the whole species dataset. The structure observed previously within the Cantal population was very weak (F_st_ = 0.05 between clusters 139R and 140R/141R) and cannot completely explain the SFS observed. In all models, the contemporary population size was as low as for the whole *A. cebennensis* dataset.

A second hypothesis was that *A. cebennensis* and *A. pedemontana* diverged after the establishment of their common ancestor to the different refugia areas, the French sub-alpine region Massif central and the alpine Piemont region in Italy. Their divergence would have resulted from the absence of gene flow due to geographical barriers. This scenario assumes a bottleneck would have happened in the common ancestor of the two species during a deglaciation period, shortly before speciation. It was tested by fitting the two-population bottleneck model 2c and comparing the results to the fitting of models 2a and 2b, which assume the divergence of the two species prior to instantaneous and exponential size change respectively. All two-populations models fitted the data very poorly based on LL values and fitting plots (Fig. S2). The lack of outgroup information and previous assumptions on any parameters of the model, the number of parameters to optimize and the potential linkage existing between our SNPs (δaδi assumes that all polymorphic sites are independent) may explain why none of the tested two-populations models fitted the data well. The fact that model 2c display the lowest LL is not in favor of a scenario including a bottleneck in the common ancestor population. The inferred divergence time between *A. cebennensis* and *A. pedemontana* was similar for models 2a and 2b (0.40 N_a_ generations ago). From model 2b we estimated that the speciation events occurred ≈ 168.000 [159676-176484] years ago.

## Discussion

As endemic species have higher probabilities to display a small census and effective population size and a highly specific ecology, reduced genetic diversity could be a limiting factor for their adaptive potential in changing environments (Ellstrand & Elam 1993; Frankham 1995; Allendorf & Ryman 2002; Leimu *et al*. 2006). In this study we evaluated the extent and structure of genetic variation in the two endemic species *A. cebennensis* and *A. pedemontana*, using samples from most of their currently known locations in southern France and northwestern Italy. Their demographic history was also investigated and their present effective population size estimated in order to determine which processes shaped the modern pattern of genetic diversity.

### Genetic structure

*Arabidopsis cebennensis* presented three clearly distinct populations corresponding to the large geographic regions Cantal, Cévennes, and Ardèche, which are represented in this study by 11, four and six samples respectively. Ardèche and Cévennes individuals were more closely related to each other. The Cantal population can be further subdivided between subpopulation 139R (4 individuals) and subpopulation 140R/141R (7 individuals). This was surprising considering that the populations 139R and 140R are geographically closer to each other (~ 4 km) than to the population 141R (~ 15 km) indicating either dispersal between the Chambeuil and the Le Siniq springs where 141R and 140R populations are respectively located, or simply shared ancestry. No reliable gene flow event could be detected between the three allopatric *A. cebennensis* populations in their recent or past history. We assume that strong barriers of gene flow exist among these units and that they are evolving independently through genetic drift. It was not surprising that an isolation-by-distance pattern (Slatkin 1993) was not apparent as only a small proportion of the total variation is expected to be spatially correlated in case of extremely limited dispersal (Meirmans 2015). In contrast, *A. pedemontana* showed only a weak clustering of individuals, probably as a result of an IBD pattern of gene flow. Dispersal is thus maintained but limited to relatively short distance within *A. pedemontana* distribution range, certainly because of the mountainous relief of the region. While *A. pedemontana* known sampling sites are all in maximum 10 km from each other, the three *A. cebennensis* populations are ~100 km away from each other with only one known additional isolated population in Aveyron, between the Cantal and the Cévennes. These long distances, the natural barriers they include, and the ecological specialization of the species explain why the connectivity among *A. cebennensis* populations is not maintained. The mean pairwise F_st_ values pointed out the isolation of these populations as they were almost as differentiated between each other than the species is differentiated from *A. pedemontana*. However, the genetic distance between *A. cebennensis* and *A. pedemontana* (0.33) was not high compared to other intra and inter-specific comparisons in the genus using neutral markers: the average pairwise F_st_ between populations of *A. halleri* from two large units separated by the Alps, was around 0.37 (Pauwels *et al*. 2012); the average F_st_ calculated between 31 *A. halleri* and 48 *A. lyrata* individuals was 0.46 ± 0.211, although this value was conservative as it was calculated from nuclear coding sequences only (Roux *et al*. 2011); the median pairwise F_st_ values calculated by Ross-Ibarra *et al*. (2008) for six natural populations of *A. lyrata* from across the range of the species were between 0 and 0.6 with 12 on 15 comparisons above 0.2. The median F_st_ between *A. cebennensis* and *A. pedemontana* was 0.22, which is equal or lower than many intra-specific comparisons within *A. lyrata*.

### Genetic diversity

Interestingly, the number and percentage of polymorphic loci were twice as high for *A. pedemontana* as for *A. cebennensis*. The nucleotide diversity (π), Watterson's estimator (θ_w_) and Nei's gene diversity (H_exp_) were all higher for *A. pedemontana* compared to *A. cebennensis*, supporting a greater genetic diversity in the Italian species despite its smaller distribution range. Only the plastid DNA variation was lower for *A. pedemontana* with only one *trnLF* plastid region haplotype while *A. cebennensis* displayed four different types (Hohmann *et al*. 2014). *A. halleri, A. arenosa*, and *A. lyrata*, the three major lineages in the genus, presented 14, 32 and 30 suprahaplotypes, much more that what was observed for the two endemics. Like *A. cebennensis* and *A. pedemontana*, *A. lyrata* and *A. halleri* are diploid, outcrossing and perennial species, but they are widely distributed. Higher average π values were also found for *A. halleri* ssp. *halleri* (0.0081) and *A. lyrata* ssp. *petraea* (0.0116) (Ramos-Onsins 2004) compared to *A. pedemontana* (0.004) and *A. cebennensis* (0.0026). The average π value in *A. thaliana* was also higher (0.0054) although the species is selfing (Nordborg *et al*. 2005). With the set of microsatellite loci included here for structure analysis, the highest number of total alleles, unique alleles and rare alleles, considering only diploids, were found within widely distributed *A. lyrata* ssp. *petraea* and *A. carpatica* (A. *arenosa* lineage; Hohmann *et al*. 2014). These diversity statistics were the lowest for *A. pedemontana* and *A. cebennensis*, as well as *A. croatica*, a third endemic species in the genus (Hohmann *et al*. 2014). Microsatellite H_exp_, calculated overall in each major lineage, is highest in *A. lyrata* (0.562 ± 0.312) and *A. arenosa* (0.560 ± 0.311), lower in *A. halleri* (0.427 ± 0.254) and much lower in *A. pedemontana* (0.259 ± 0.167) and *A. cebennensis* (0.189 ± 0.130). When calculating mean H_exp_ over SNPs only (and not over the total number of mapped sites), we obtained similar values with 0.33 for *A. cebennensis* and 0.23 for *A. pedemontana*. Only for this comparison *A. pedemontana* does not display a higher genetic diversity compared to *A. cebennnensis*. Overall, we observe that the genetic variation level in the two endemic species is lower than for the other *Arabidopsis* species with wide distribution range and the same mating system (self-incompatibility). For comparison, the narrow endemic *Aquilegia thalictrifolia*, distributed in a few valleys of the Italian South-Eastern Alps, also displayed a higher average H_exp_ (0.68 ± 0.19), which is twice as high as for the two *Arabidopsis* endemics (Lega *et al*. 2014). In *A. cebennensis* π, θw and H_exp_ were low and almost similar for the three subpopulations, and similar to what was estimated for the whole species. A similar level of genetic diversity is partitioned among and within the subpopulations in this species. It shows that gene flow was too limited to ensure that variation is shared over the whole distribution of the species, and led to the strong population structure observed. Overall, both *A. pedemontana* and *A. cebennensis* display low levels of genetic variation at the taxon and population levels, as would be predicted based on their narrow geographic range.

### Comparison of population genetics inference methods

The SNPs from the GBS analysis and the microsatellites markers gave similar results with regard to population structure, despite the differences in sampling size. In *A. cebennensis*, the level of admixture observed between the Ardèche and Cévennes populations with the microsatellite data confirm that these two populations had more recent genetic exchanges than with the Cantal populations. Concerning genetic diversity statistics, the GBS analyses gave much lower H_exp_ values than the microsatellites markers when both polymorphic and non-polymorphic sites were accounted for. As it takes in account the number of SNPs detected over the total number of sites, this approach gives a more realistic estimation of the level of heterozygosity over the whole genome. Nonetheless the two methods of polymorphism detection gave similar results in terms of ratio of H_exp_ between the two endemic species. The genome-wide SNPs and the multiple microsatellite loci were thus equally useful in inferring population structure and comparing the level of diversity between related species in our study. One advantage of using GBS is to screen thousands of polymorphism that are subject to the full range of evolutionary processes acting across the genome (mutation, drift, selection) (Narum *et al*. 2013) and improve the precision of demographic inferences by greatly increasing the number of putatively neutral markers assayed.

### Past decline of the endemics populations

Our overall neighbor-net network with *A. lyrata, A. cebennensis* and *A. pedemontana* confirmed previous observations that the two endemic species, although clearly separated, are sister species (Koch & Matschinger 2007; Hohmann *et al*. 2014). Additionally, 10% and 5% of the total polymorphism in *A. cebennensis* and *A. pedemontana* are shared polymorphisms, confirming the joint evolutionary history of the two species. As the two species are ecologically, morphologically and in certain case phylogenetically closer to *A. halleri* (Hohmann *et al*. 2014), we assumed that they diverged from a common ancestor related to A. *halleri* lineage. *A. halleri* is distributed in the whole alpine chain while the endemic species occur in refuge areas west of the mountainous range, at the highest altitude in the surroundings. The periphery of European Alps, in particular the south east and the west of the mountain range, harbors many small refugia with hundreds of endemic plant species, which result from Pleistocene glaciation cycles (Comes & Kadereit 2003; Tribsch & Schonswetter 2003; Schönswetter *et al*. 2005). We therefore hypothesized that the evolutionary histories of both species have been influenced by Pleistocene climate oscillations: the two cold-adapted species diverged from their common ancestor in the western alps during a glaciation period; with the next deglaciation phases the populations migrated or became restricted to refugia in higher altitude where they could survive the warming temperatures, resulting in a strong overall population size decrease; because of their adaptation to their specific refugia habitat and/or competition, they could not expand back to their unknown original distribution range, forming relictual populations.

The demographic analysis confirmed that both endemic species have undergone a strong decline in the past and did not recover to the present days, although we were not able to state if this decline happened through a strong bottleneck or through exponential population size decrease. The population decline was particularly strong for *A. cebennensis*, which displayed a contemporary effective population size *N_e_* of 140 to 450 individuals, only 0.1% of the size of its ancestral population. *A. pedemontana's* decline was less drastic, as the species exhibits a modern *N_e_* of 50,000-55,000 individuals, half of its ancestral population size. In comparison, Roux *et al*. (2011) estimated *N_e_* of ~82,000 and ~79,200 for the two more widespread relatives *A. halleri* and *A. lyrata*. According to our estimates, *A. pedemontana* population started decreasing 18,000 to 41,000 years ago (ya), while *A. cebennensis'* decline was much more recent, i.e. 200 to 3,700 ya. The time of divergence of the two species was estimated at ~170,000 ya. However these estimates are doubtful for two reasons: (i) the two-populations demographic models within which the time of divergence parameter was optimized fitted the observed dataset poorly; (ii) the time values were converted in units of years by using an average generation time of two years. Because the two species are long-lived perennials, propagate vastly vegetatively, and their dispersal could be limited for some times by their specific ecology and the space competition observed in their habitat (pers. obs.), the chosen generation time as well as the divergence time and times of decline could be underestimated. The divergence of *A. cebennensis* and *A. pedemontana* falls in the medium Pleistocene (Ionien), which corroborates our hypothesis. These results fit with the time of radiation calculated for all *Arabidopsis* lineages with n = 8 chromosomes (all except *A. thaliana)* (1.63 Mya; Hohmann *et al*. 2015), and for *A. halleri* ssp. *halleri* (~335,000 ya; Roux *et al*. 2011). While *A. pedemontana's* decline falls still in the late Pleistocene, the decline of *A. cebennensis* seems to have occured in modern time (Holocene). As the demographic models (1a and 1b) used to estimate this value did not fit *A. cebennensis* observed data SFS perfectly, and *A. cebennensis* population structure was not included in the models, we can not say with certainty that this very recent estimate is accurate. Our separate demographic analysis of *A. cebennensis* Cantal population showed that the strong population structure of *A. cebennensis* alone does not explain completely the right-shifted site frequency spectrum observed for this species and the extremely low population size that was estimated. While population structure could explain the excess of intermediate frequency alleles, a recent strong bottleneck could explain the lack of rare alleles (Luikart *et al*. 1998) and the small population size. In the future, the evolutionary history of the three *A. cebennensis* populations will be further investigated with an extended and more representative sampling scheme, which will also allow us to integrate the population structure into the demographic analysis. Particularly we will test if the structure and the sampling size together have created a false bottleneck signal for this species (Chikhi *et al*. 2010).

### Consequence on adaptive potential

In summary we described two endemics dissimilar patterns of genetic variation and differentiation. Due to a potentially rapid and recent decline, and its split into three allopatric populations, *A. cebennensis* currently exhibits a very low effective population size and low levels of genetic diversity. Although the absence of gene flow between these three sub-populations initiated their divergence by genetic drift or divergent selection (according to F_st_ values), the genetic diversity among them is also low (according to π values). The probability of finding strongly differentiated genotypes and phenotypes in the different geographic region where the species occurs is consequently small. *Arabidopsis pedemontana* population also declined rapidly in the past but not as strongly as for *A. cebennensis*, and *A. pedemontana* effective population size is still relatively high. Distance-limited dispersal is ongoing within its distribution range but did not contribute to create much diversity between the locations. Indeed the overall diversity level remains low. The contemporary population size, levels of gene flow versus genetic drift and demographic history of both species can explain why we observed higher level of diversity for *A. pedemontana* compared to *A. cebennensis*, despite its smaller distribution range. A historically larger population size in *A. pedemontana* could also have contributed to its higher level of genetic diversity.

These first results are not optimistic regarding the adaptive potential of the two species in case of environmental change. The relatively recent and on-going global climate warming could impact the habitats of these two species by two means: 1) In Europe, it was already observed that climate change causes a general upward migration of plants to higher altitudes where they may compete with locally adapted endemic plants (Lenoir *et al*. 2008); 2) The increasing drought that could result from rising temperatures and decreasing precipitation predicted to be induced by climate change could particularly impact the highly specific riverine habitat of the two species. As a result, the occurrence of the two species in high altitude habitats, where they can not escape the increased competition, and their habitat specialization render them particularly vulnerable to climate change (Gottfried *et al*. 2012). Predicting populations persistence during environmental change is complex as it depends on numerous factors (Reed *et al*. 2011). However, with the relatively low levels of genetic diversity observed, we can predict that the two *Arabidopsis* endemic species may experience reduced relative adaptive ability and a higher risk of genetic extinction. The risk could be relatively higher for *A. cebennensis*, as small population size decrease the rates of adaptive evolution (Strasburg *et al*. 2011; Lanfear *et al*. 2014) and do not allow the populations to sustain further decline before evolutionary rescue (Gonzalez *et al*. 2012).

## Acknowledgments

We thank Elizabeth Kokai-Kota for laboratory assistance, and Markus Stetter and Fabian Freund for help with the population genetic analysis. Financial support was provided to K.J.Schmid by the DGF Grant SCHM1354/6-1 within the DFG priority programme 'Adaptomics'.

## Data Accessibility

‐ GBS raw reads: Genbank NCBI submission SRP072277
‐ Biallelic SNPs final vcf files: DataDryad
‐ Demographic analysis scripts: Supporting information

## Author Contributions

JJ coordinated the project, performed the GBS sequencing and population genetic analysis and wrote the manuscript. NH and MAK conducted the microsatellite and plastid DNA sequences analysis and contributed to write the manuscript. MB contributed to the sample collection and DNA extraction. AS helped with the sample collection. TM assisted with the GBS reads processing and population genetic analysis. KJS designed and coordinated the project and contributed in writing the manuscript. All authors read and approved the final manuscript.

## Appendices

Figure S1. Barcode sequences and design of double-digest GBS libraries

Figure S2. Comparison of the fitted demographic models. The optimized parameters and log likelihood values (LL) are indicated for each model. The site frequency spectrum is plotted for each model in red and compared to the observed data SFS plotted in blue. Ac = *Arabidopsis cebennensis;* Ap = *A. pedemontana*.

Figure S3. Genetic assignments of *A. cebennensis* and *A. pedemontana* populations from population structure analysis with microsatellite and plastid DNA sequences: A) Summary of microsatellite analysis; B) Distribution of *A. cebennensis* populations analyzed for microsatellite variation. The color-code refers to STRUCTURE results shown with C; C) STRUCTURE plot of the microsatellite data with K=3 (highest probability) for 144 *A. cebennensis* individuals from 10 populations. The chloroplast genome type are indicated for each population below the graph (T, BG, A, BH); D) Spatial distribution of *A. pedemontana* populations analyzed for microsatellite variation. The color-code refers to STRUCTURE results shown with E; E) STRUCTURE plot of the microsatellite data with K=2 (highest probability) for 40 *A. pedemontana* individuals from 9 different localities. All individuals carried the same single chloroplast genome type (A).

Figure S4. Neighbor Net network constructed for *A. cebennensis, A pedemontana* and outgroup *A. lyrata*. Bootstrap supports are indicated on the branches.

Figure S5. Cross validation values for each iteration and number of clusters in ADMIXTURE analyses: A) *A. cebennensis;* B) *A. pedemontana*.

Figure S6. TreeMix Maximum likelihood tree of *A. cebennensis* three large populations with *A. pedemontana* as outgroup.

Figure S7. Manhattan plots of F_st_ values between: A) *A. cebennensis* and *A. pedemontana* individuals; B) the three large populations of *A. cebennensis;* along *A. lyrata* scaffolds used as reference sequence.

Figure S8. Manhattan plots of nucleotide diversity values (π) within: A) all *A. cebennensis* individuals; B) all *A. pedemontana individuals;* along *A. lyrata* scaffolds used as reference sequences.

Table S1. Q-estimates (admixture proportion at the individual level, expressed as the fraction of each population that contribute to the individual genome) for *A. pedemontana* individuals, with two populations assumed in ADMIXTURE analysis. Individuals for which more than 10% of each population contribute to the genome are highlighted in grey.

Table S2. SNPs filtering after SNP-calling analysis: 1) Number of total sites, variant sites and polymorphic sites at the successive steps of the filtering process, 2) Mean read depth per site, 3) Percentage of variant and polymorphic sites on the total number of sites; when calculating the % of polymorphic loci, only intraspecific variants and SNPs were taken in account; '>50' = only SNPs with data for at least 50% of the sampled individuals; '>70' = only SNPs with data for at least 70% of the sampled individuals.

Table S3. SNP-calling results for subsets of 21 randomly chosen *A. pedemontana* samples: numbers of intraspecific biallelic SNPs with data for at least 70% (>70) of the sampled individuals are indicated.

Text S1. Detailed Materials and Methods

## Supporting information

Supporting information 1. Identification, location and collection informations for samples of *A. cebennensis* and *A. pedemontana* used in the GBS, microsatellite and plastid DNA analysis.

Supporting information 2. Summary of GBS results in number of raw and processed reads overall and per individual. The shaded rows in the second table point out the individuals with less than 5,0SNPs in the final calling, which were removed from subsequent analysis.

Supporting information 3. Python script implementing all models used in the demographic analyses.

